# Fungal hyphae colonization by *Bacillus subtilis* relies on biofilm matrix components

**DOI:** 10.1101/722272

**Authors:** Bodil Kjeldgaard, Stevanus A. Listian, Valliyammai Ramaswamhi, Anne Richter, Heiko T. Kiesewalter, Ákos T. Kovács

**Author notes:** Contributed equally (Bodil Kjeldgaard, Stevanus A. Listian).

## Abstract

Bacteria interact with their environment including microbes and higher eukaryotes. The ability of bacteria and fungi to affect each other are defined by various chemical, physical and biological factors. During physical association, bacterial cells can directly attach and settle on the hyphae of various fungal species. Such colonization of mycelia was proposed to be dependent on biofilm formation by the bacteria, but the essentiality of the biofilm matrix was not represented before. Here, we demonstrate that secreted biofilm matrix components of the soil-dwelling bacterium, *Bacillus subtilis* are essential for the establishment of a dense bacterial population on the hyphae of the filamentous black mold fungus, *Aspergillus niger* and the basidiomycete mushroom, *Agaricus bisporus*. We further illustrate that these matrix components can be shared among various mutants highlighting the community shaping impact of biofilm formers on bacteria-fungi interactions.

## 1. Introduction

Biofilm development by a plethora of bacteria and fungi has been pragmatically studied separately in the laboratory. Biofilm formation is abundant in the environment [1] and these biofilm communities likely comprise both bacteria and fungi in addition to higher eukaryotes grazing or residing these habitats. Interaction among bacteria and fungi extends physical associations, and furthermore includes chemical communications, ranging from antibiosis and metabolic exchange to specific signaling and chemotaxis [2]. These direct or indirect interactions among bacteria and fungi alters the physiology, growth, movement, differentiation, pathogenesis, reproduction and/or survival of either or both partners [2]. During physical interaction between bacteria and fungi, stable association is possibly facilitated by the production of a viscous biofilm matrix. Indeed, the production of a biofilm matrix has been suggested for various bacterial isolates on the hyphae of the ectomycorrhizal fungus, *Laccaria bicolor* [3], however, the essentiality of the biofilm matrix in long term attachment has not been confirmed before. Several molecular determinants have been described to be important for bacteria-fungi interactions, including motility, quorum sensing, bacterial secretion system, secondary metabolites (see an extended list reviewed in [4]). Here, we examined the role of biofilm components of *Bacillus subtilis* during association with filamentous Ascomycota and Basidiomycota. *B. subtilis* is a soil-derived Gram-positive bacterium and long-studied model for physiology, genetics and differentiation, including motility, sporulation, biofilm formation [5]. Phenotypic diversification of this bacterium is influenced by intertwinement of global regulators, including Spo0A that has been originally described to be involved in production of heat and pressure resistant spores [6]. In addition, Spo0A has been later shown to impact protease secretion, toxin production, and biofilm development, thus involved in regulation of genes related to cell fate decision of the bacterium [7]. Formation of biofilm in *B. subtilis* requires a complex matrix composed of exopolysaccharide (EPS), TasA amyloid fiber, and a surface localized hydrophobin, BslA [8–13]. The former two matrix components are essential for the establishment of a floating biofilm on the air-medium interface (referred to as pellicle), the creation of complex vein-like structures on agar medium, or attachment to the root surface of plants [8–10,14,15]. BslA protein creates a shield around the pellicle and colony biofilm to avoid penetration of hydrophobic fluids and its deletion alters microstructure of colonies [12,13,16], but it is not indispensable for establishment of floating biofilms [17,18]. The ability and degree of biofilm matrix production correlates with attachment to the plant root surface in hypotonic cultures and also with the ability of the bacterial strains to protect tomato plant against wilt disease caused by *Ralstonia solanacearum* [10,19]. As the matrix components are secreted from the cells, mutants lacking either EPS or TasA are able to complement each other and create a functional biofilm both *in vitro* and on the plant root [10,11,20].

A previous study demonstrated that attachment of *B. subtilis* to the mycelia of *Aspergillus niger* leads to metabolic alterations in both partners [21]. Microarray experiments revealed that genes responsible for single cell motility in *B. subtilis* are downregulated 3 hours after attachment [21], suggesting settlement of the bacterial cells on the fungal hyphae. However, transcriptions of biofilm-related genes were not altered at such an early stage of interaction compared to planktonic cells.

Here, we revisited the interaction of *B. subtilis* with *A. niger* dissecting the co-cultures after 24 hours of incubation. We demonstrate that long-term attachment depends on the genes required for exopolysaccharide and amyloid fiber production, but neither the synthesis of hydrophobin nor the ability for single cell motility are required for the bacterial settlement on the fungal mycelium. Moreover, the secreted matrix components can complement single gene deletion mutants. Finally, we show that such a matrix-dependent colonization of fungal hyphae is not only restricted to *A. niger*, but similarly required for biofilm formation on the mycelia of the basidiomycete, *Agaricus bisporus*.

## 2. Material and methods

### 2.1 Strains, media composition and culturing conditions

All the bacterial and fungal strains used in this study are listed in Table 1. Bacterial strains were cultivated in Lysogeny Broth medium (LB-Lennox, Carl Roth, Germany; 10 g l^-1^ tryptone, 5 g l^-1^ yeast extract and 5 g l^-1^ NaCl) supplemented with 1.5 % Bacto agar if required. Supplemented LB medium was prepared by adding 1 % 1 mol l^-1^ MgSO_4_ and 0.1 % 0.1 mol l^-1^ MnCl_2_ to basic LB medium. *A. niger* was cultivated on potato dextrose glucose agar (PDA) medium (Carl Roth, Germany; potato infusion 4 g l^-1^, glucose 20 g l^-1^, agar 15 g l^-1^, pH value 5.2 ± 0.2) to harvest spores and in LB medium for formation of macro-colonies. *A. bisporus* was cultivated on PDA plates or potato dextran glucose broth (PDB) medium (Carl Roth, Germany; potato infusion 4 g l^-1^, glucose 20 g l^-1^, pH value 5.2 ± 0.2). TB526 was obtained using natural competence [22] by transforming genomic DNA extracted from NRS2097 to TB34 and selecting for chloramphenicol resistance, followed by verifying the mutation by PCR. To select for resistant bacterial colonies after transformation, 5 μg l^-1^ chloramphenicol was used.

**Table 1.** Bacterial and fungal strains used in this study. Cm^R^, Spec^R^, Km^R^, and Tet^R^ denote chloramphenicol, spectinomycin, kanamycin, and tetracycline resistance cassettes, respectively.

| **Bacterial strains** |  |  |
| --- | --- | --- |
| *B. subtilis* strains | Characteristics | Reference |
| DK1042 | NCBI3610 *comI*<sup>Q12I</sup> | [31] |
| TB34 | DK1042 *amyE*::P<sub>hyperspank</sub>-*gfp*, Cm<sup>R</sup> | [32] |
| TB35 | DK1042 *amyE*::P<sub>hyperspank</sub>-*mKATE2*, Cm<sup>R</sup> | [33] |
| TB36 | DK1042 Δ*hag*::Km<sup>R</sup>; *amyE*::P<sub>hyperspank</sub>-*gfp*, Cm<sup>R</sup> | [34] |
| TB37 | DK1042 Δ*hag*::Km<sup>R</sup>; *amyE*::P<sub>hyperspank</sub>-*mKATE2*, Cm<sup>R</sup> | [34] |
| TB421 | DK1042 Δ*spo0A*::Km<sup>R</sup>; *amyE*::P<sub>hyperspank</sub>-*gfp*, Cm<sup>R</sup> | [32] |
| TB524 | DK1042 Δ*eps*::Tet<sup>R</sup>; *amyE*::P<sub>hyperspank</sub>-*gfp*, Spec<sup>R</sup> | [11] |
| TB525 | DK1042 Δ*eps*::Tet<sup>R</sup>; *amyE*::P<sub>hyperspank</sub>-*mKATE2*, Spec<sup>R</sup> | [11] |
| NRS2097 | NCIB3610 Δ*bslA*::Cm<sup>R</sup> | [35] |
| TB526 | DK1042 Δ*bslA*::Cm<sup>R</sup>; *amyE*::P<sub>hyperspank</sub>-*gfp*, Spec<sup>R</sup> | This study |
| TB538 | DK1042 Δ*tasA*::Km<sup>R</sup>; *amyE*::P<sub>hyperspank</sub>-*gfp*, Spec<sup>R</sup> | [11] |
| TB539 | DK1042 Δ*tasA*::Km<sup>R</sup>; *amyE*::P<sub>hyperspank</sub>-*mKATE2*, Spec<sup>R</sup> | [11] |
| **Fungal strains** |  |  |
| Species name | Strain and collection numbers | Reference |
| *A. bisporus* | H39 / CBS 122262 | [36] |
| *A. niger* | N402 / ATCC 64974 / CBS 132248 | [37] |

### 2.2 Hyphal colonization assay of A. niger

Spores of *A. niger* cultures grown at 28°C for 7-12 days on PDA plates were harvested using 10 ml saline tween solution (8 g l^-1^ NaCl and 0.05 ml l^-1^ Tween 80) and filtered through Miracloth (Millipore; Billerica, USA) following the protocol described in [21]. The spore solution was centrifuged 5000 rpm for 10 min and resuspended in saline tween solution. Spores were stored at 4°C until use, but no more than 14 days. Around 3*10^5^ spores ml^-1^ were inoculated into 25 ml of LB medium and shaken at 120 rpm at 28°C for 24 hours. 4-5 macro-colonies were transferred into a single well of a 24-well plate and culture was supplemented with 0.01 mol l^-1^ MgSO_4_ and 0.1 mmol l^-1^ MnCl_2_ as indicated for supplemented LB medium. 10 µl of *B. subtilis* overnight culture grown at 37°C (single or mixed culture) was inoculated into each well. The co-cultures were incubated at 28°C with shaking at 170 rpm for 24 h. Following incubation, the pellicle biofilm formed at the air-liquid interface was removed from each well. Subsequently, *A. niger* macro-colonies were washed two times with sterile MilliQ water before imaging.

### 2.3 Hyphal colonization assay of A. bisporus

Three-week old *A. bisporus* culture, grown on PDA plates at 25°C was wetted with 10 ml of physiological saline and scraped thoroughly with a spreader. Afterwards, 1 ml of the hyphal suspension was incubated in 25 ml PDB at 25°C without agitation for 8-14 days. After eight days, *A. bisporus* macro-colonies were floating within the media. One macro-colony each was transferred into a well of a 24-well microtiter plate. Remaining PDB was removed and the macro-colonies were washed once with physiological saline. 1 ml of supplemented LB medium was added to each macro-colony and inoculated with 10 µl of *B. subtilis* overnight culture grown at 37°C (single or mixed culture). The microtiter plate was incubated at 25°C with shaking at 170 rpm for 22-24 h. Subsequently, *A. bisporus* macro-colonies were washed three times with physiological saline.

### 2.4 Sample preparation and confocal laser scanning microscopy (CLSM)

Washed fungal macro-colonies were transferred to microscope slides and gently sealed with cover slips. Fungal hyphae colonization was analysed with two confocal laser scanning microscopes (LSM 710 (Carl Zeiss) or TCS SP8 (Leica) both equipped with an argon laser and a Plan-Apochromat 63x/1.4 Oil objective). Fluorescent reporter excitation was performed at 488 nm for green fluorescence and at 561 nm for red fluorescence, while the emitted fluorescence was recorded at 540/40 nm and 668/86 nm (Zeiss) or 520/23 nm and 700/90 nm (Leica) for GFP and mKATE, respectively, (wavelength/bandwidth). For generating multi-layer images, Z-stack series with 1 μm steps were acquired and processed with the software ImageJ (National Institutes of Health). Maximum intensity was used to merge five chosen stacks of the green fluorescent, red fluorescent and overlay channel. Merging of the five stacks was done to integrate signals from the bacterial cells at different focus planes. The bright field channel is represented by only one brightness-adjusted image from one of the stacks used to obtain the fluorescent images.

## 3. Results and discussion

### 3.1 Colonization of the A. niger hyphae by B. subtilis depends on the global regulator, Spo0A

Transcriptome analysis of attaching *B. subtilis* cells on *A. niger* mycelia has revealed that expression of genes related to single cell motility is reduced compared to non-attached cells [21] suggesting that the bacterial cells switch from planktonic to sessile state of growth. The global transcriptional regulator, Spo0A influences hundreds of genes in *B. subtilis* that determine bacterial cell fate, e.g. differentiation into swimming, biofilm forming, or sporulating cell types [7]. Therefore, the impact of *spo0A* gene deletion in *B. subtilis* was first assayed during colonization of the hyphae of *A. niger*. As expected, bacterial cells of the *spo0A* strain showed reduced hyphal colonization compared to the wild type and only planktonic cells were observed around *A. niger* (Fig. 1 A,B). Driven by the observation that colonization of plant root by *B. subtilis* requires secreted biofilm matrix components [10,19], we have inspected if addition of wild type strain restores attachment of *spo0A* mutant cells on the fungal hyphae. Indeed, co-inoculation of wild type and *spo0A* cells in 1:10 ratio rescued attachment of *spo0A* strain to the hyphae, suggesting that secreted factors by the wild type strains are sufficient for the establishment of biofilm by the two strains on the fungal mycelia (Fig. 1C). Alternatively, signaling molecules produced by the wild type strain, but absent in the *spo0A* mutant could induce the molecular factors responsible for hyphal biofilms.

**Fig. 1.**
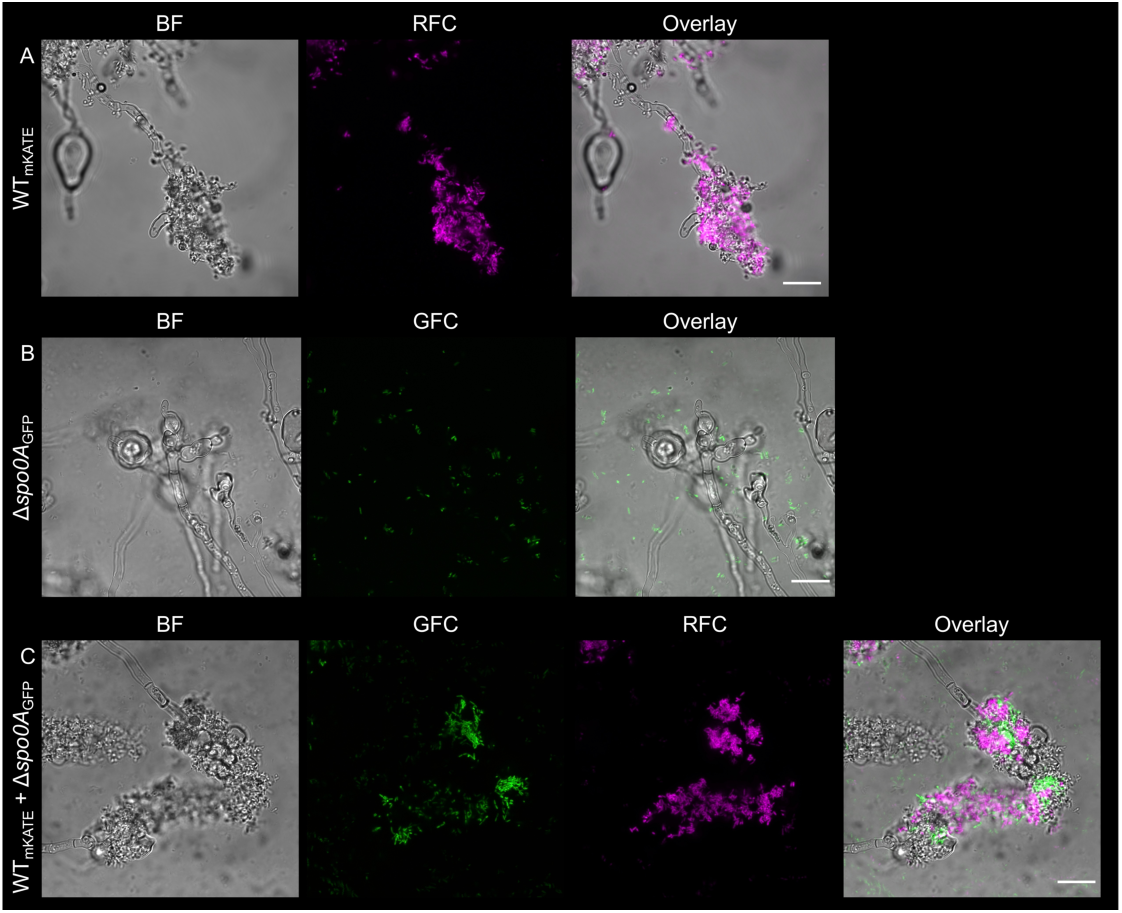
Colonization of the *A. niger* hyphae by *B. subtilis* (A) wild type, (B) *spo0A* mutant and (C) the co-culture of 10:1 GFP-labeled *spo0A* mutant and mKATE-labeled wild type. GFC: green fluorescent channel; RFC: red fluorescent channel. Scale bars indicate 20 μm. The images presented are representative examples selected from independent samples repeated on different days.

### 3.2 Hyphal colonization depends on B. subtilis biofilm matrix components, EPS and TasA

Biofilm formation of *B. subtilis* depends on various secreted components, including EPS, TasA and BslA. The transcription of these operons involved in the production of matrix is indirectly dependent on Spo0A. Therefore, we have tested how single deletion of *eps* operon, *tasA* or *bslA* genes impacts fungal colonization by *B. subtilis*. Unlike the *bslA* mutant, removal of either EPS or TasA hindered the bacterial biofilm development on the hyphae of *A. niger* (Fig. 2A-C). This suggests that both core components of the matrix, EPS and TasA contribute to establishment of stable biofilm on the mycelia, while the hydrophobin BslA is not necessary for stable attachment. In accordance, previous studies found that BslA is not essential for the establishment of floating pellicle biofilm of *B. subtilis*, but is required for its repellency and the microstructure of colonies [12,13,17,18]. Finally, gene deletion in the flagellin coding gene was assayed for biofilm establishment during mycelia colonization. Although motility is not essential for generation of biofilms, it has been described to be critical for fast formation of *B. subtilis* biofilm [23]. Cells lacking single cell motility could colonize the hyphae of *A. niger* (Fig. 2D), similar to the wild type strain (Fig. 1A). The importance of bacterial flagella and type 4 pili was previously described during co-migration of *Burkholderia terrae* with fungal hyphae through soil [24], suggesting that essentiality of bacterial motility for fungal interaction could depend on the environment, bacterial physiology or specific properties of the fungal and bacterial cell surfaces. In addition, other examples highlight that fungal hyphae could facilitate bacterial swimming along the hyphae and therefore niche colonization [25,26]. In our simple laboratory system, the bacterium-fungus co-cultures are mildly agitated, which allows firm contact of the bacterial cells and the mycelia. Under these conditions, motility is plausibly not essential.

**Fig. 2.**
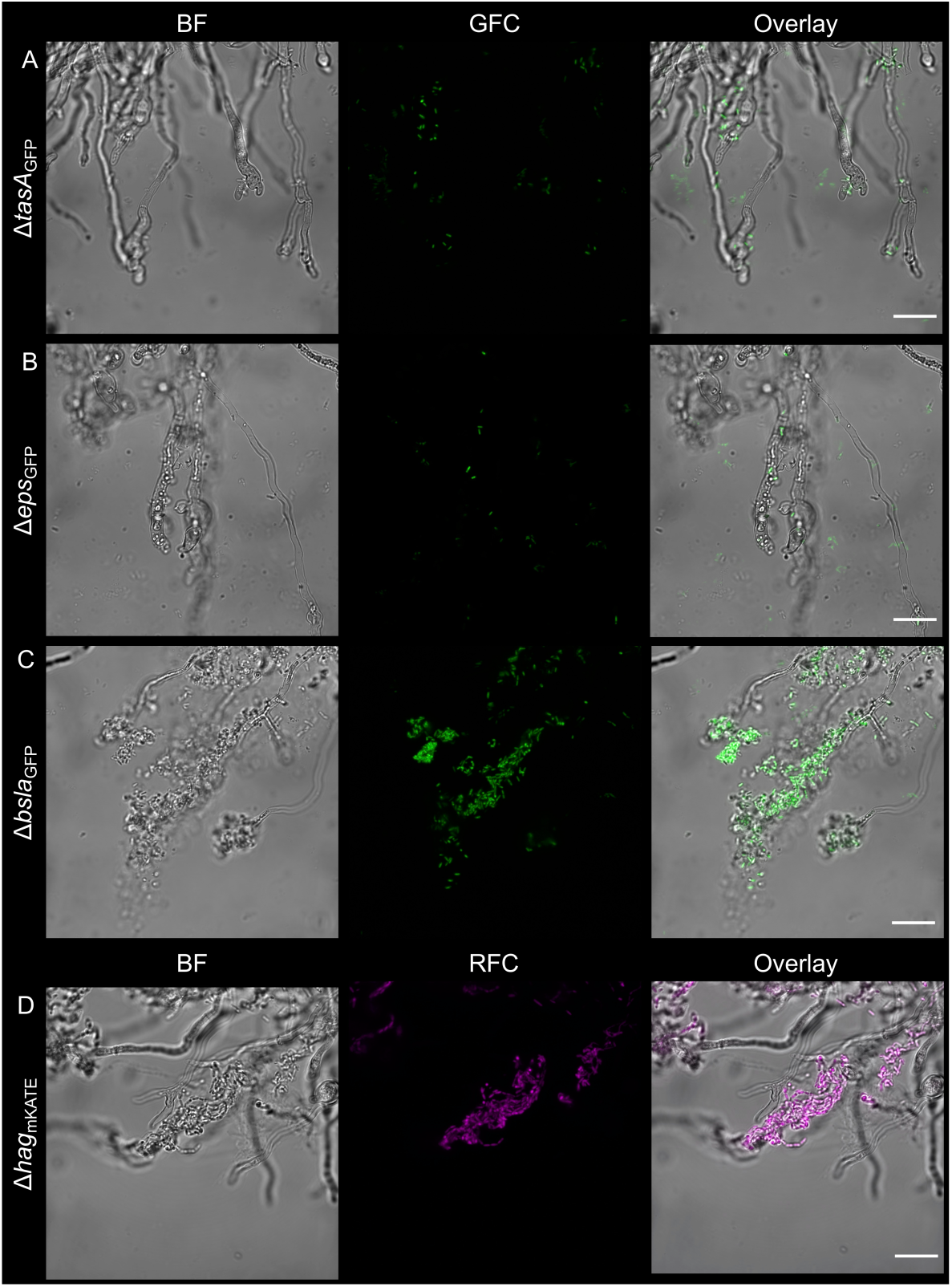
*B. subtilis* lacking biofilm components, (A) TasA or (B) EPS are unable to grow on the *A. niger* hyphae, while strains lacking (C) the production of hydrophobin, BslA or (D) motility established stable colonization on the mycelia. GFC: green fluorescent channel; RFC: red fluorescent channel. Scale bars indicate 20 μm. The images presented are representative examples selected from independent samples repeated on different days.

### 3.3 Secreted biofilm matrix components are shared among mutants

To complement the mutations and lack of biofilm formation ability of the bacterial strains on the hyphae of *A. niger*, various co-cultivations were tested. Deficiency in EPS or TasA production was complemented either by co-cultivation of mutant strains together or by addition of the wild type strain to biofilm matrix mutant *B. subtilis*, irrespective of fluorescent labeling (Fig. 3). Previous studies demonstrated that secreted biofilm matrix components complement pellicle biofilm formation [20,27] as well as plant colonization [10,11]. Notably, mixing *eps* and *tasA* mutant *B. subtilis* strains results in genetic division of labor, thus increased population productivity both during pellicle biofilm formation at the air-liquid interface and during plant root colonization [11,28].

**Fig. 3.**
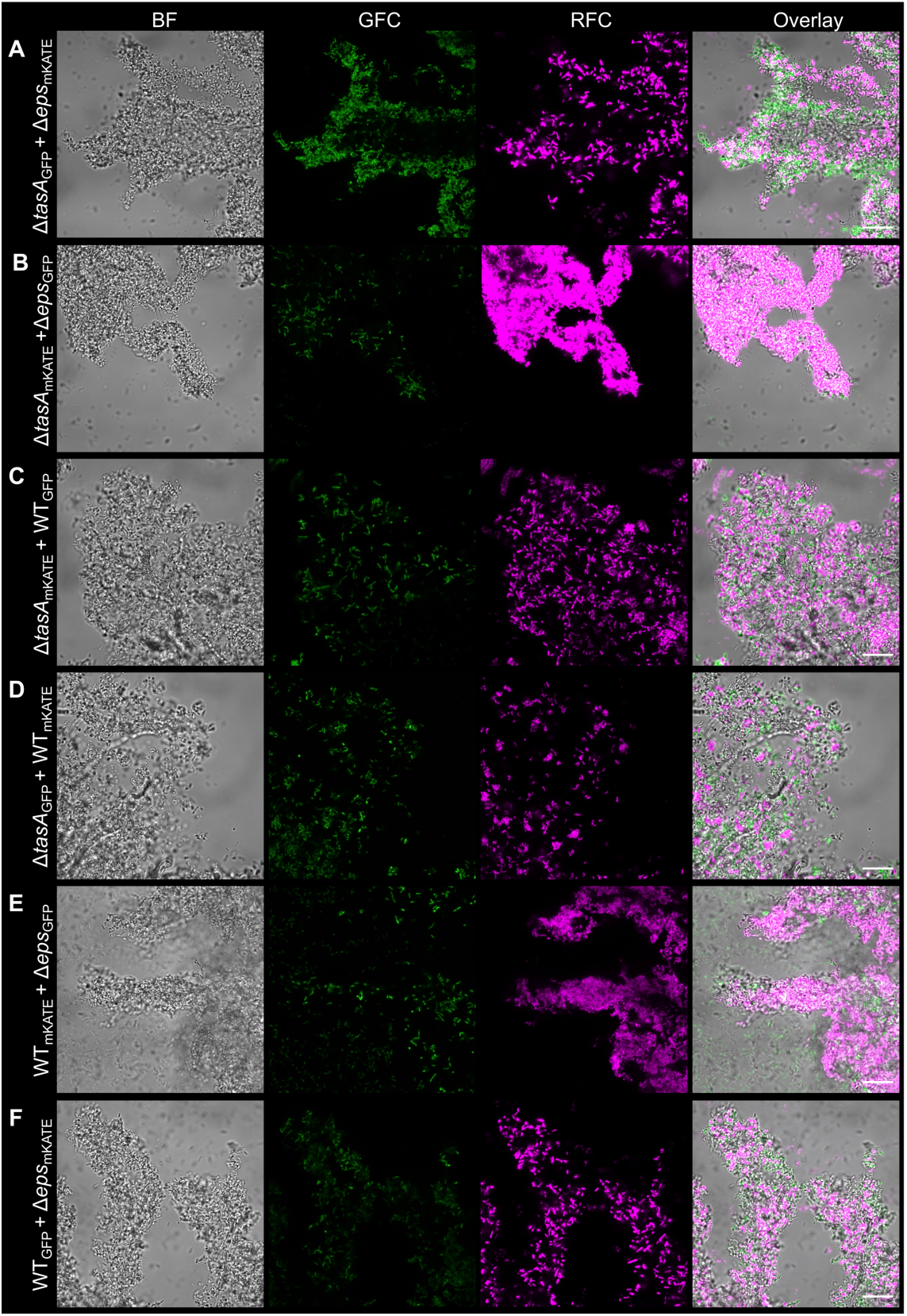
Mutations in *tasA* and *eps* genes were complemented by the co-inoculation of the mutants (A and B), and by the wild type strain (C-F) including a fluorescent marker swap, resulting in fungal colonization by the mutants. GFC: green fluorescent channel; RFC: red fluorescent channel. Scale bars indicate 20 μm. The images presented are representative examples selected from independent samples repeated on different days.

### 3.4 Biofilm formation of B. subtilis on the basidiomycete mycelia

The above described simple co-culture system comprises a bacterium, *B. subtilis* and an Ascomycota fungus, *A. niger* that can be both isolated from soil, however, their co-occurrence in nature has never been reported according to our knowledge. Therefore, we examined fungus-attached biofilm formation by *B. subtilis* on a more ecologically relevant host. *In vitro* laboratory experiments suggest that *B. subtilis* cells can colonize the hyphae of the ectomycorrhizal fungus, *L. bicolor* [3] and the basidiomycete *Coprinopsis cinerea* [29]. In addition, *B. subtilis* has been isolated from the soil directly underneath a troop of growing *Paxillus involutus* mushrooms [30] and below an *Agaricus* sp. fruiting body (Kiesewalter and Kovács, unpublished observation). Therefore, we set out to examine the importance of biofilm matrix components during mycelia colonization of button mushroom, *A. bisporus*. In agreement with the above observations on *A. niger* from this study, hyphal biofilm formation by *B. subtilis* was diminished by deletion of *spo0A, epsA-O*, or *tasA* genes, while removal of *bslA* or *hag* genes did not impact colonization properties (Fig. 4).

**Fig. 4.**
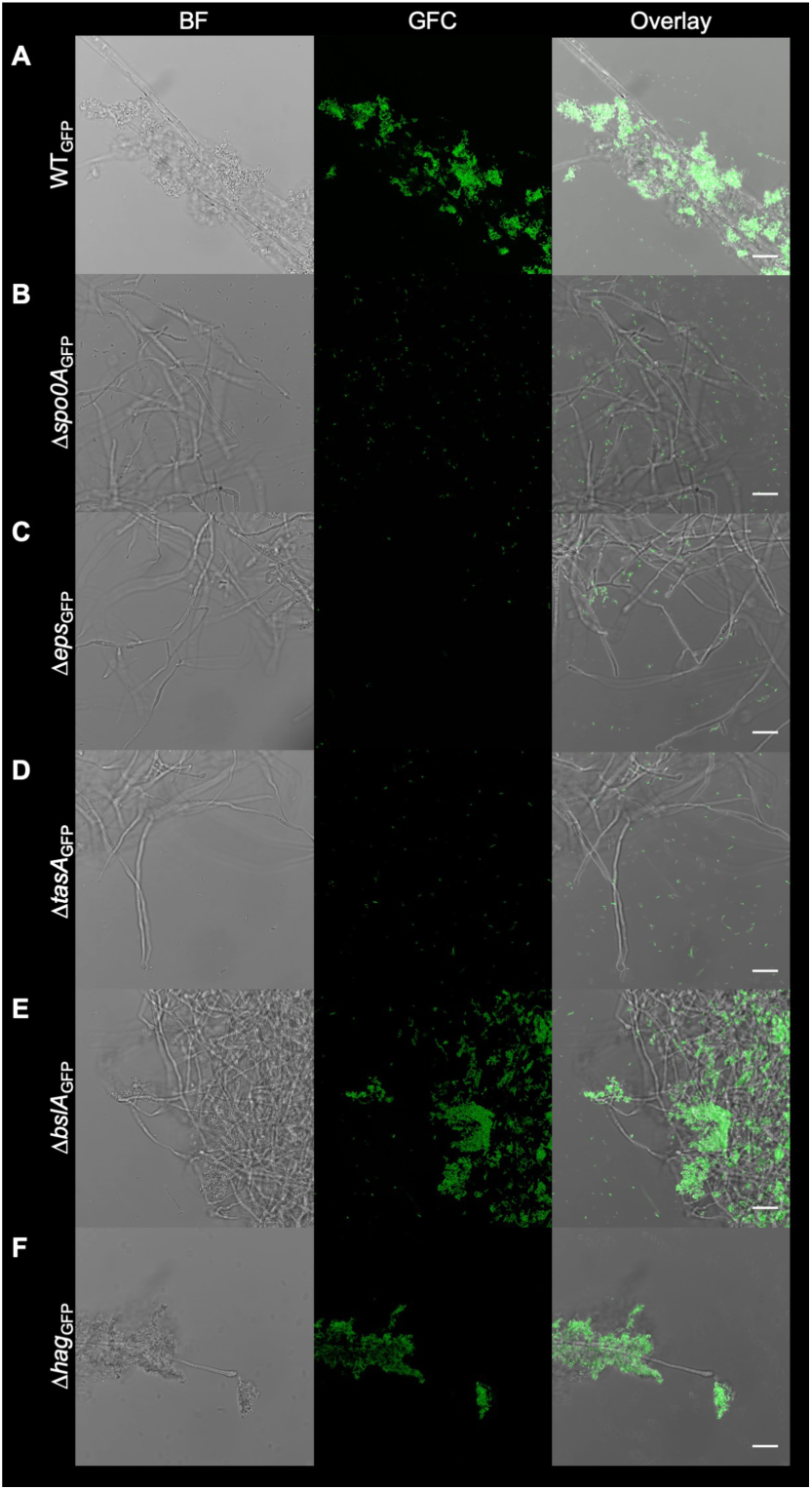
Attachment of *B. subtilis* on *A. bisporus* hyphae. *B. subtilis* wild type (A) and its corresponding mutants were co-cultivated with *A. bisporus* macro-colonies. Fungal macro-colonies were subsequently washed and visualized with CLSM. The images were recorded at the edge of the fungal macro-colonies. GFC: green fluorescent channel. Scale bars represent 20 µm. The images presented are representative examples selected from different positions of the fungal macro-colonies on independent samples repeated on different days.

Further, complementation of the matrix mutants, *eps* and *tasA* strains was performed using a co-culture of the mutants or by addition of wild type to the mutants (Fig. 5). The experiments with the basidiomycete, *A. bisporus* strengthens the observation that fungal hyphae colonization and biofilm formation by *B. subtilis* depends on secreted biofilm components. This suggest that production of a matrix by biofilm-proficient bacterial species in nature could potentially facilitate the establishment of multi-species biofilms on fungal hyphae. Indeed, the presence of biofilm matrix and extracellular DNA has been demonstrated for a numerous bacterial species during colonization of ectomycorrhizal fungi, including *L. bicolor* [3]. Our genetic approach further supports the importance of bacterial biofilm matrix production during long term colonization of fungal mycelia.

**Fig. 5.**
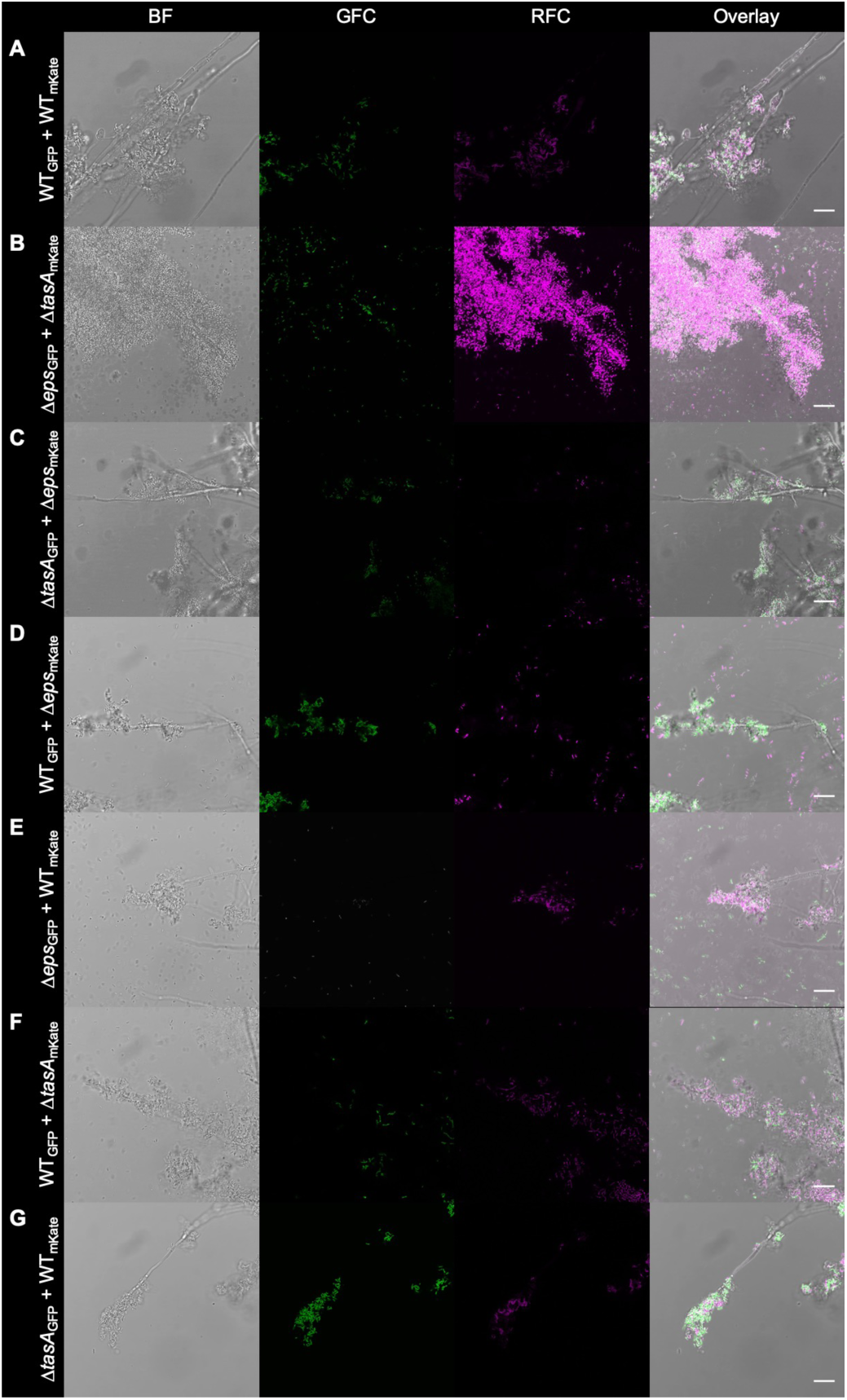
Attachment of mixed-culture *B. subtilis* strains to *A. bisporus* hyphae. Overnight cultures of different *B. subtilis* strains were mixed and co-cultivated with *A. bisporus* macro-colonies. Fungal macro-colonies were subsequently washed and visualized with CLSM; pictures were taken approximately near the center of the macro-colonies. GFC: green fluorescent channel; RFC: red fluorescent channel. Scale bars represent 20 µm. The images presented are representative examples selected from different positions on independent samples.

## 4. Conclusion

Bacterial biofilms in the laboratory have been mostly studied using inert substrates, however, during infections or in the environment, bacteria also interact with eukaryotes, including their hosts (from humans and animals to plants) or the co-habitants of their milieu. Here, we demonstrate that biofilm matrix components of *B. subtilis* are essential for colonization of the hyphae of *A. niger* and *A. bisporus*. In addition, the secretion of these matrix components is sufficient to rescue biofilm formation of matrix deficient strains suggesting that social interaction likely shapes the co-evolution of fungi and bacteria in the environment. This leads to the appearance of specific interactions, including primary metabolite cross-feeding, molecular recognition, and potential induction of secondary metabolite production.

## Acknowledgement

We thank Nicola Stanley-Wall for the comments on our bioRxiv submission.

